# Deep mapping of the TCR-antigen interface using pMHC-pseudotyped viruses and yeast display

**DOI:** 10.1101/2025.08.25.671738

**Authors:** Stephanie A. Gaglione, Jonathan Krog, Frances R. Keer, Emma N. Finburgh, Elif Kulaksizoglu, Connor S Dobson, David Gifford, Aziz Al’Khafaji, Paul C. Blainey, Michael E. Birnbaum

## Abstract

T cell receptor (TCR) specificity is central to the efficacy of T cell therapies, yet scalable methods to map how TCR sequences shape antigen recognition remain limited. To address this, we introduce VelociRAPTR, a library-on-library approach that combines yeast-displayed TCR libraries with pMHC-displaying virus-like particles (pMHC-VLPs) to rapidly screen millions of TCR-antigen interactions. We show that pMHC-VLPs efficiently bind TCRs on yeast and generate equivalent data to recombinantly produced pMHC protein. We then apply VelociRAPTR to screen 47 million variants of the A6 and 868 TCRs against 92 pMHCs simultaneously, mutating both the CDR3 loops and cognate peptides. The resulting CDR3-pMHC maps reveal biased recognition patterns, where mutations to CDR3 loops can selectively constrain or broaden specificity to peptide analogs. These insights provide a foundation for engineering TCRs with defined pMHC binding profiles and improving models that predict TCR-antigen interactions, including the prediction of off-target recognition. By coupling the scale of yeast display with the modularity of VLPs, VelociRAPTR offers a generalizable strategy for generating deep, high-throughput protein-protein interaction data.

## Introduction

T cells mediate adaptive immunity by recognizing specific antigens via the T cell receptor (TCR). Most T cells express αβ TCRs, which bind short peptide antigens displayed by major histocompatibility complex (MHC) proteins. Contact between the TCR and peptide-MHC (pMHC) is primarily driven by the TCR’s complementarity-determining region (CDR) loops. The hypervariable CDR3α and CDR3β loops are the most diverse and mediate contact with the peptide, while the less diverse CDR1 and CDR2 loops primarily interact with the MHC helices^1–3^. TCRs must balance the specificity required for precise antigen recognition with sufficient cross-reactivity to surveil a large landscape of potential antigens^4–7^ with a TCR repertoire limited to 10^8^ unique clones per individual^8,9^.

Beyond their role in natural immunity, TCRs are increasingly being explored as therapeutic modalities in cancer, infection, and autoimmunity, with recent FDA approvals including a TCR- based bispecific engager for uveal melanoma^10^, a tumor-infiltrating lymphocyte therapy for melanoma^11^, and an engineered TCR-T therapy for synovial sarcoma^12^. TCRs are attractive therapeutically due to their ability to sensitively target intracellularly derived antigens presented by MHCs. The ability to characterize, and potentially optimize, TCR affinity for a pMHC is therefore of considerable therapeutic interest. Nonetheless, reliably mapping TCRs to cognate antigens and comprehensively characterizing TCR cross-reactivity remain challenging due to the low affinity of TCR-antigen interactions and expansive scale of potential binding pairs^13^.

Despite recent advances^14–22^, current approaches to study and enhance TCR-antigen interactions remain limited in scale. Directed evolution-based methods have been used to increase the low affinity of TCRs while maintaining antigen specificity. These methods, including phage^23^ and yeast^23–27^ display, generate large libraries of TCR mutants, primarily targeting the CDR3 regions for mutagenesis, followed by iterative rounds of selection against an antigen of interest^25,28–34^. Although capable of high-throughput TCR generation and screening, these assays are limited by the need to produce and test each antigen as a soluble protein individually. Conversely, methods exist to screen vast antigen libraries against a single TCR using yeast^6^ or mammalian cell display^16^, but these approaches cannot screen many TCRs simultaneously.

Our group and others have worked to address these limitations by developing library-on-library screening approaches for the TCR-pMHC interaction. These methods, including pMHCs displayed on engineered lentiviruses being used to barcode TCR-pMHC interactions by binding to the TCR or enabling TCR-mediated infection of T cells^14,15,17,35^, and TCR-pMHC interaction pairing via engineered yeast mating^36^, show promise in simultaneously assessing diverse TCRs and antigens. However, these approaches remain limited by their overall scale, difficulties conducting rounds of selection, and inefficiencies in capturing low-affinity TCR-antigen interactions.

To overcome these constraints, we aimed to pair the scalability of yeast display with the utility of lentiviral display. Here, we introduce VelociRAPTR, an approach for generating deep library-on-library protein-protein interaction data. Using libraries of fluorescent pMHC-displaying virus-like particles, we screen millions of yeast-displayed TCR variants against hundreds of pMHCs simultaneously over iterative rounds of enrichment. Enriched TCRs and pMHCs are then identified by bulk or single-cell sequencing. We apply this method to systematically interrogate two TCR model systems, screening 47 million CDR3α/β-mutated TCRs against 92 pMHCs simultaneously to investigate how altered CDR3s shape TCR cross-reactivity and to identify key residues for TCR- antigen recognition. We envision this approach as a means of rigorously evaluating candidate therapeutic TCRs for off-target binding and reactivity^37–39^ and generating training data for deep learning models of TCR recognition^40–45^.

## Results

### pMHC-pseudotyped VLPs bind TCR-displaying yeast

Viruses and virus-like particles (VLPs) are well-precedented for displaying proteins of interest in the context of gene delivery^46,47^, vaccines^48^, mammalian cell screens^14,17^. VLPs can display membrane-bound proteins, encode nucleic acid barcodes, be generated in high-throughput, and be used to examine TCR-antigen interactions in primary T cells using ENTER-seq and V- CARMA^14,35^. Extending this, we reasoned that VLPs could synergize with yeast display for massive library-on-library screens. By staining yeast libraries with pooled, barcoded pMHC-VLPs and iteratively sorting, we can enrich TCR-binding populations. The identities of TCR-pMHC binding pairs can then be identified by bulk or single-cell sequencing of the yeast-VLP complexes.

To test whether pMHC-VLPs bind TCR-expressing yeast in an antigen-dependent manner, we examined the A6 and 868 model system TCRs. The A6 TCR recognizes the HTLV-1 peptide Tax (LLFGYPVYV) bound to HLA-A2^49^ and has been thoroughly characterized structurally and biochemically^26,50–53^. Similarly, the 868 TCR targets the HIV-1 peptide SL9 (SLYNTVATL) bound to HLA-A2^54,55^ with high affinity. Both TCRs stably express on yeast as single-chain variable fragments (scTv), composed of variable TCR domains connected by a flexible linker in a Vβ- linker-Vα format^26^.

As a proof-of-concept, we stained yeast expressing A6 and 868 TCRs with fluorescent Tax and SL9 pMHC-VLPs. Mimicking recombinantly produced pMHC tetramer, we engineered VLPs to display pMHCs as single-chain trimers of covalently linked peptide, MHC, and beta-2- microglobulin (β2M) and encoded a barcode corresponding to pMHC identity in the viral genome (**Figure 1a**). To identify pMHC-VLPs bound to yeast, we incorporated an HIV gag-fluorophore fusion protein into the viral capsid^56^. We observed that pMHC-VLPs selectively stain yeast expressing their cognate TCRs, with SL9 VLPs binding 868 TCR^+^ yeast and Tax VLPs binding A6 TCR^+^ yeast (**Figure 1b**). In addition to measuring VLP fluorescence directly, VLP signal can be amplified with a secondary anti-β2M antibody. Confirming that this approach is compatible with lower affinity TCRs, we generated and tested VLPs displaying pMHCs known to bind the A6 TCR across a range of physiologically relevant affinities: 0.9 μM (Tax)^52^, 7 μM (Tel1p549-557, MLWGYLQYV)^50^, 41 μM (Tax-V7R, LLFGYPRYV)^52^, and 123 μM (HuD87-95, LLYGFVNYI)^57^ (Supplementary Figure 1).

**Figure 1.**
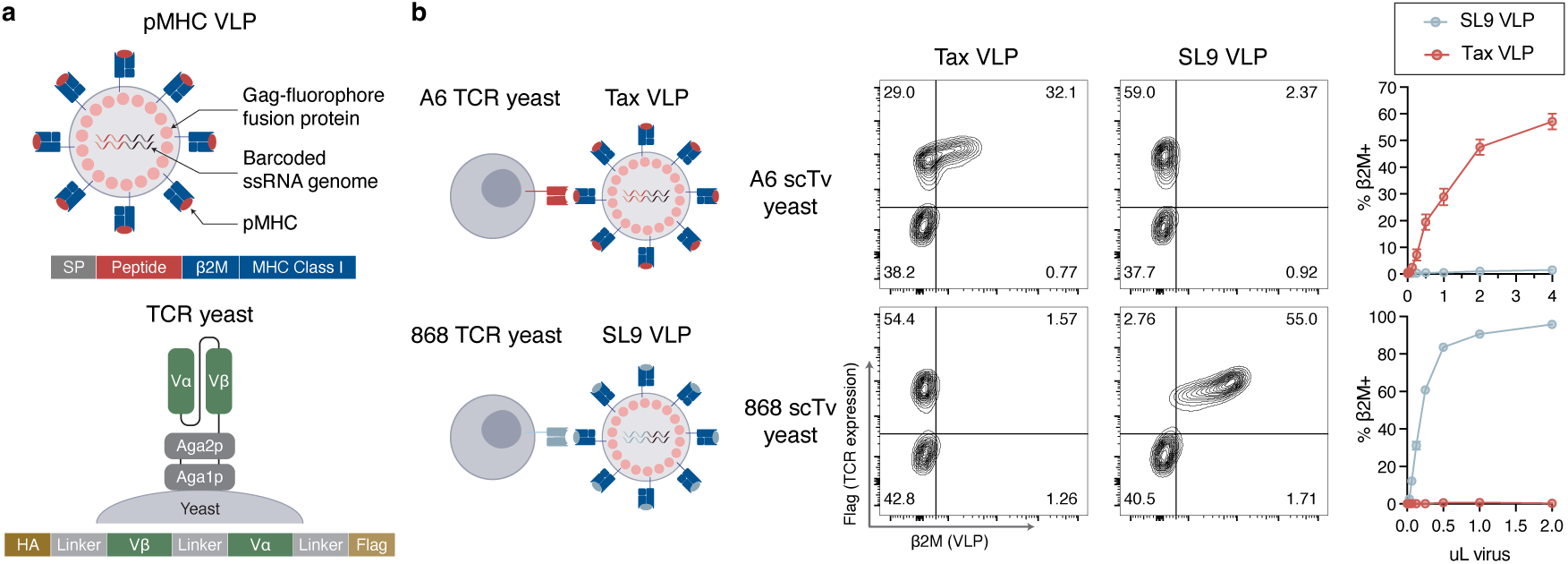
**pMHC-pseudotyped VLPs bind antigen-specific yeast**. **a**, Schematic of pMHC-VLP and TCR- expressing yeast. pMHC-pseudotyped virus-like particles (VLPs) display pMHCs as disulfide-stabilized single- chain trimers (peptide-β2M-A2) and incorporate fluorophores conjugated to Gag and a barcoded ssRNA viral genome. The A6 TCR is expressed as a single-chain TCR (scTv fusion) consisting of two linked variable regions (VH and VL)^26^. The alpha and beta variable domains are connected by a flexible linker in the orientation Vβ-L- Vα at the C-terminus of the yeast cell wall protein Aga2. **b,** Specific binding of A6- and 868-TCR yeast by pMHC-VLPs displaying cognate antigens Tax and SL9, respectively, with minimal off-target binding. Flow plots depict pMHC binding of yeast expressing a linked epitope tag (Flag), including a secondary stain of VLP-bound yeast with anti-β2M. Line plots depict dose-response of VLP staining. Plots show mean ± S.D. of 3 technical replicates.

### pMHC-VLPs enrich antigen-specific CDR3s equivalently to recombinant pMHC

Following validation that pMHC-VLPs bind TCR-displaying yeast in an antigen-specific manner, we next assessed whether pMHC-VLPs perform comparably to recombinantly produced pMHCs in a screening context. As an initial test, we constructed a library of ∼6,800 A6 TCR variants by randomizing three CDR3α positions (D98, S99, W100) with close peptide contact as determined by the previously resolved A6-Tax/HLA-A2 co-crystal structure^53^ (**Figure 2a**).

**Figure 2.**
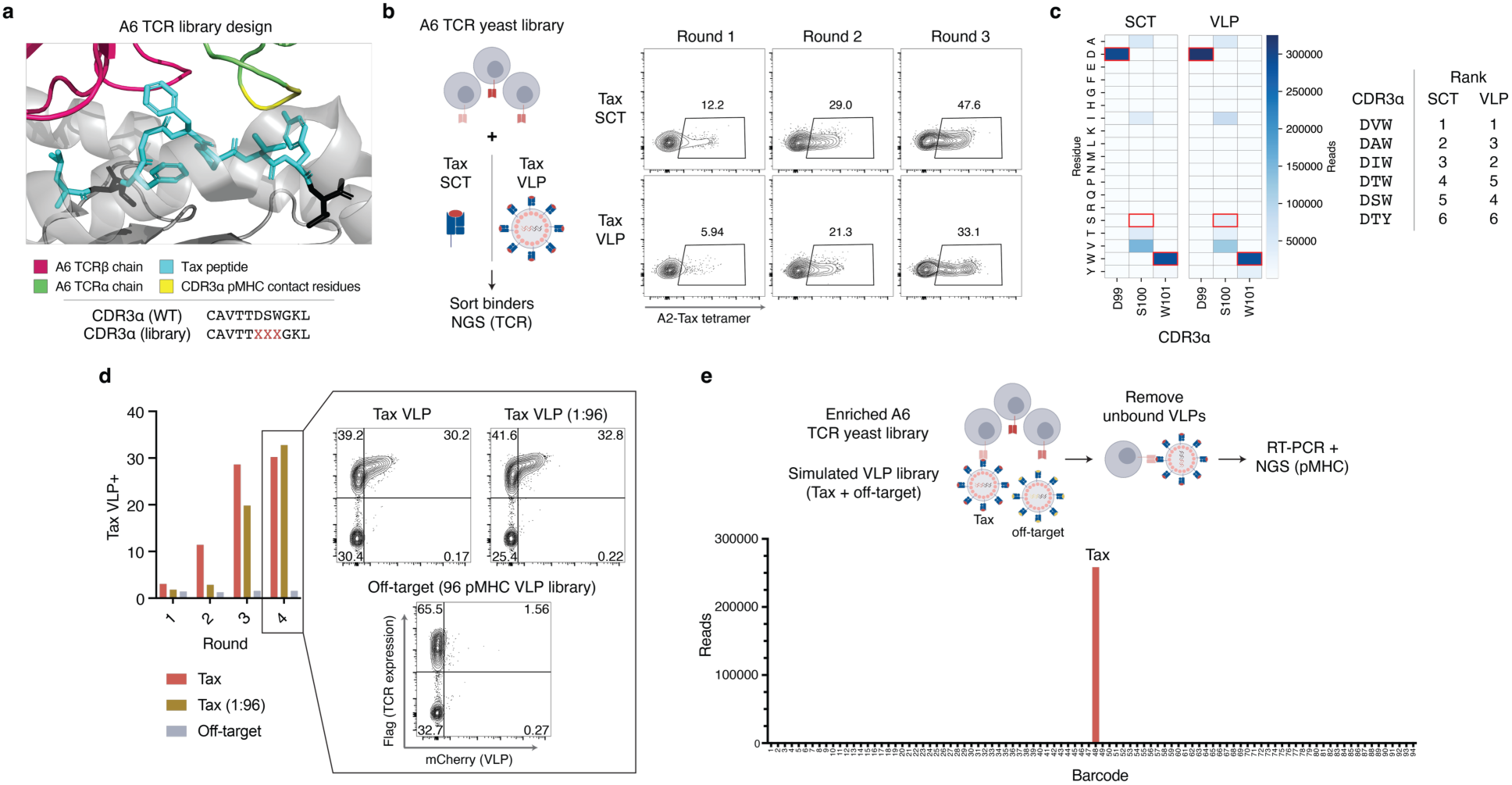
pMHC-VLPs enrich A6 CDR3α variants comparably to recombinant pMHC. **a**, Design of A6 TCR variant library, mutating three residues in CDR3α (D99, S100, W101), indicated by X, with close peptide contact. **b**, Comparison of A6 TCR variant library enrichment with multimerized recombinantly produced Tax single-chain trimer (SCT) and Tax VLP. Flow plots depict yeast after each of three rounds of selection, stained with Tax tetramer. **c,** Heatmaps depicting reads per residue at each CDR3α position after three rounds of selection with recombinant SCT and VLP. Table lists the relative rank of CDR3α sequences in enriched yeast. **d**, Library-scale enrichment of naïve A6 TCR library with Tax VLPs (red), an off-target 96 pMHC-VLP pool (grey), and a proportional mixture of Tax VLPs in 96 pMHC-VLPs (gold). Bar plot depicts VLP staining after each round and flow plots show staining after round 4. **e,** Reads per pMHC barcode following staining of enriched yeast with 97 pMHC-VLP pool where Tax is the only on-target pMHC.

To benchmark pMHC-VLPs against the current standard, yeast libraries were iteratively stained and sorted with both pMHC-VLPs and recombinantly expressed, multimerized pMHC (**Figure 2b**). After each of three rounds of selection, we assessed the library enrichment via next-generation sequencing (NGS). Of the ∼6800 potential CDR3α variants, the top six enriched sequences were identical between recombinant protein-based selection and VLPs and enriched residues at each position are virtually identical (**Figure 2c**). In addition to the wild-type CDR3α (DSW), the top three enriched CDR3α sequences conserve D98 and W100 with aliphatic residues at position 99 (S99V, S99A, S99I).

Compared with recombinantly produced protein, barcoded VLPs enable highly multiplexed screening and the ability to capture pMHC identity via NGS. To test multiplexing capacity, we next screened the A6 TCR variant library against multiple antigens simultaneously. Complementing Tax pMHC-VLPs, we generated a 96-member pMHC-VLP library consisting of peptides derived from common viruses, none of which are expected to enrich A6 TCR variants^15,17^. We then screened the A6 CDR3α variant TCR library against three conditions: Tax VLPs alone, the 96 off-target VLP library, and a 97-member VLP library combining on-target Tax VLPs with 96 off-target VLPs at a 1:96 ratio. The off-target 96-VLP library failed to enrich yeast while the Tax-only and 97-VLP library enriched A6 TCR variants (**Figure 2d**). The absence of enrichment by the off-target VLP library confirms the minimal nonspecific binding between pMHC-VLPs and TCR-displaying yeast and confirms the well-characterized specificity of the A6 TCR^49^.

Following validation that a simulated library of pMHC-VLPs can enrich TCR variants, we amplified and bulk sequenced pMHC barcodes from VLPs interacting with selected TCRs. Staining the enriched A6 variant library with a pool of 97 pMHC-VLPs, including the cognate Tax pMHC-VLP, revealed that over 99% of reads from bound VLPs correspond to Tax (**Figure 2e**). These results demonstrate that a ∼100-member pMHC-VLP library specifically enriches TCR variants and that both TCRs and pMHC identities can be efficiently captured via NGS.

### Library-on-library screens of CDR3α/β variant libraries against peptide mimotopes

While prior approaches have enabled deep mutational scanning of either TCRs or peptides in isolation^23–27,33^, few simultaneously interrogate both sides of the recognition interface. Mapping these high-dimensional interaction networks is essential to efforts modeling the molecular basis of TCR specificity and cross-reactivity^58–62^. Building on our proof-of-concept screen, we applied our library-on-library VLP-yeast system to rapidly screen TCR variants against a library of peptide mimotopes to identify CDR3 motifs with varied cross-reactivity (**Figure 3a**). We expanded the scale of the A6 TCR variant library by randomizing 3 positions with close peptide contact in both CDR3α and CDR3β^53^ (CDR3α positions D98, S99, and W100 and CDR3β positions L99, A100, G101), increasing the library diversity from ∼6800 to 47 million TCR variants. Concurrently, we generated a library representing 47 million variants of the 868 TCR, also randomizing 3 positions in both CDR3s with close contacts based upon an X-ray crystal structure^55^ (CDR3α positions N94, S95, Y97 and CDR3β positions T96, V97, 101G). To screen both TCR libraries for reactivity, we constructed a VLP library consisting of single amino acid substitutions of known key TCR contact residues for Tax and SL9 (37 variants of Tax, 37 variants of SL9), in addition to 18 off-target pMHCs representing common HLA-A2-binding viral epitopes **(Figure 3b and Table S1**). Collectively, 47 million TCR variants and 92 pMHCs represent 4.3 billion possible interacting pairs.

**Figure 3.**
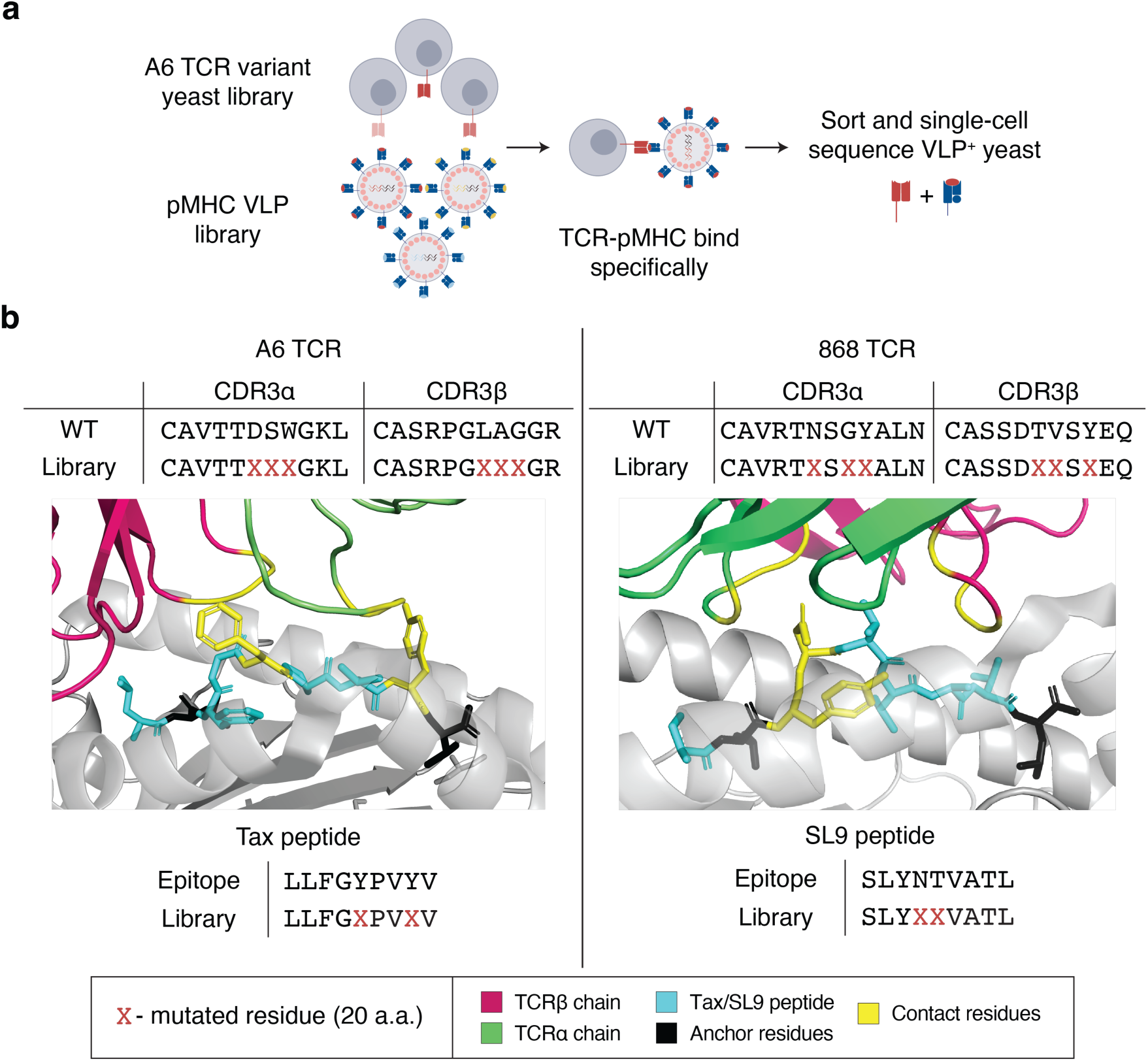
Yeast and VLP library construction for large-scale assay. a, Schematic showing co-incubation of yeast-displayed TCR libraries with pMHC-displaying VLP libraries. Pairs are enriched over rounds of selection with sorting and sequencing of VLP-stained yeast. **b,** Design of A6 and 868 TCR variant libraries, varying both at 3 positions in the CDR3, denoted by X. The corresponding 92-VLP library includes 37 variants of each cognate peptide (Tax and SL9), varied at two positions denoted by X.

To facilitate capture of low-frequency TCRs, we optimized a magnetic-assisted cell sorting (MACS) protocol to easily enrich rare VLP-labelled yeast. After staining the large yeast variant library with 92 VLPs in pool, we completed a secondary stain with a PE-conjugated anti-β2M antibody followed by incubation with anti-PE magnetic microbeads. Bead-based enrichments of VLP-labeled yeast were able to isolate VLP-bound yeast with both high efficiency and purity (**Supplementary** Figure 2).

We iteratively screened the A6 and 868 variant TCR libraries against the 92 pMHC-VLP library and compared the VLP library against single on-target pMHC-VLPs (Tax or SL9) as controls. Both the individual VLPs displaying the cognate peptide for each TCR and the 92 pMHC-VLP library enriched strongly (**Figure 4a**). We next sequenced enriched yeast via NGS to examine and compare CDR3α/β clonotypes enriched by the VLP library and wild-type pMHC-VLPs. Both wild-type Tax and the VLP library strongly enrich CDR3α residues at positions 99 and 101, consisting of Aspα99, Trpα101, and Tyrα101, and CDR3β residues at positions 98 and 100, specifically Lysϕ398, Methϕ398, and Trpϕ3100 (**Figure 4b**). However, for both CDR3α and CDR3β, wild-type Tax clearly enriches specific hydrophobic residues at position 100 (CDR3α) and 99 (CDR3β) compared with the VLP library (i.e. Alaα98, Isoα98, Methϕ399, Trpϕ399), suggesting that the broader set of pMHC variants binds a more diverse set of TCRs.

**Figure 4.**
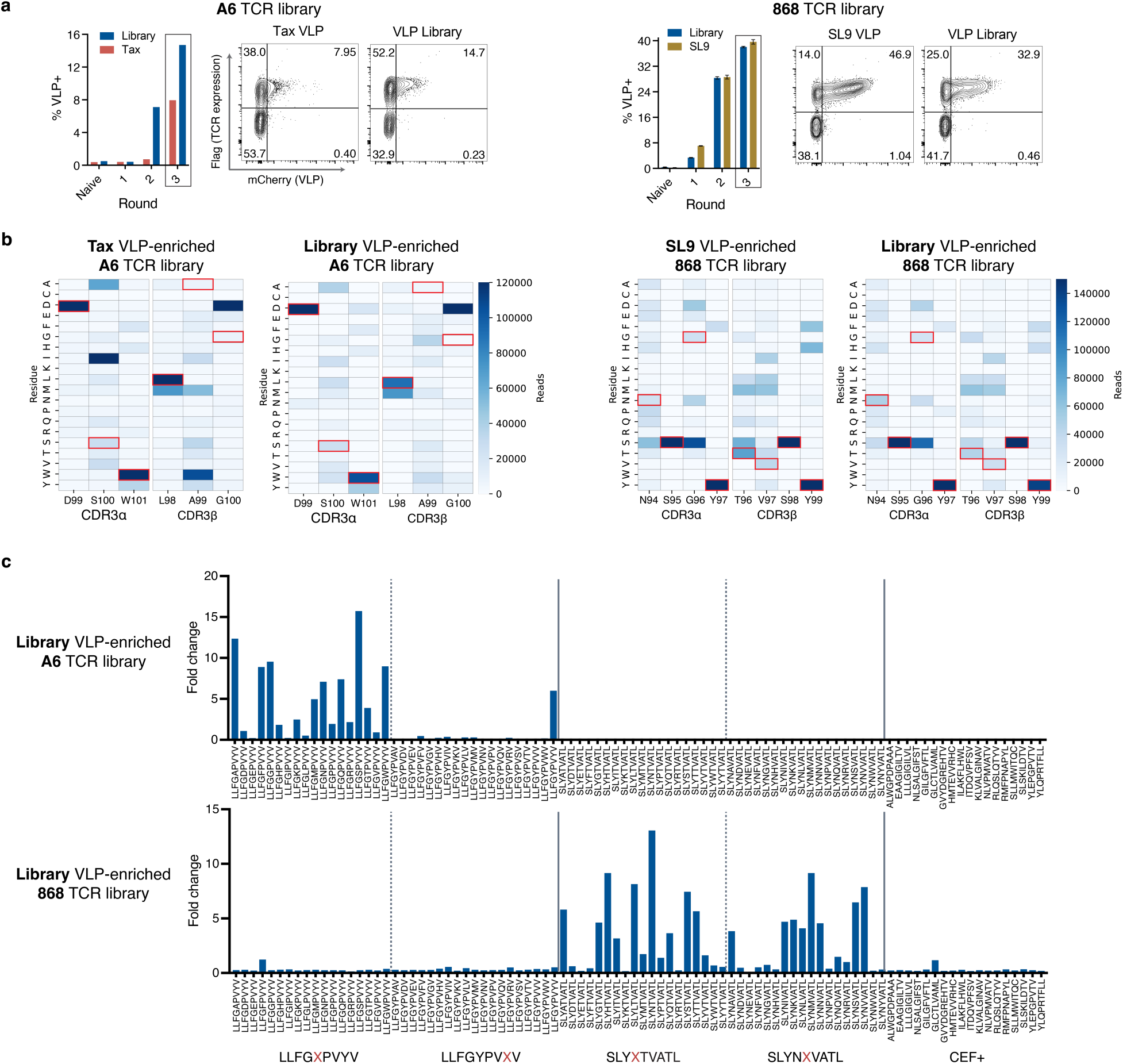
VelociRAPTR enables library-on-library enrichment of A6 and 868 TCR variants. **a**, Enrichment of A6 and 868 CDR3α-CDR3β TCR variant libraries against Tax-A2, SL9-A2, and a 92-pMHC library containing 37 Tax mimotopes, 37 SL9 mimotopes, and viral A2-binding antigens. Bar plot depicts staining of yeast after each round and flow plots show VLP staining after the third round. **b,** Heatmaps depicting reads per residue at each mutated CDR3α and CDR3β position for both A6 and 868 TCR libraries after three rounds of enrichment. Data shown for yeast enriched with a single pMHC-VLP and the VLP library. **c**, NGS data of barcoded pMHC-VLPs bound to enriched A6 and 868 TCR libraries. Fold change is calculated as the proportion of reads per VLP relative to the proportion of reads for each VLP in the 92-VLP library. Bars represent mean ± S.D. of 3 technical replicates.

In contrast, the 868 TCR preferences for SL9 closely matched the VLP library at all positions. The 868 TCR tolerates more diverse residues within CDR3α and CDR3β, except for Serα96, Tyrα97, and Tyrϕ399. Demonstrating the specificity of this pMHC-VLP binding, we separately sequenced VLPs bound to enriched yeast for each TCR library via NGS (**Figure 4c**). Each yeast-displayed TCR library enriched for peptides resembling its cognate antigen, with little signal observed for the variants of the other TCR’s antigen or for the off-target pMHC controls. Notably, A6 variants tolerate epitopes with alternate residues at position 5 (LLFG<u>X</u>PVYV) but not position 8 (LLFGYPV<u>X</u>V). 868 variants bind SL9 variants more promiscuously, with some low detected binding of off-target peptides. To confirm that a 92-member pMHC-VLP library accurately reflects individual pMHC binding, we generated individual VLPs for each Tax variant and stained enriched A6 TCR yeast separately with each VLP, demonstrating that individual pMHC-VLP staining aligns with bulk sequencing data (**Supplementary** Figure 3).

### Sequencing yeast-VLP pairs to match CDR3s to pMHCs

With two TCR variant libraries enriched against libraries of pMHC variant VLPs, we next sought to map interacting TCR-antigen pairs by sequencing both the single-chain TCRs from yeast (CDR3α and CDR3β) and pMHC barcodes from VLPs. Existing methods like single-cell RNA sequencing have been used to decode TCR-pMHC pairs in viral-mammalian display systems^14,17^. We therefore adapted a single-cell sequencing approach to be compatible with yeast to identify which CDR3α and CDR3β transcripts associated with pMHC-VLP barcodes. We additionally used the NGS data from the yeast enrichments (**Figure 4b**) to identify top paired CDR3α/β clonotypes. Using this approach, we extracted top CDR3α-pMHC and CDR3β-pMHC pairs for both A6 and 868 TCRs, focusing on the A6 TCR for subsequent validation (**Figure 5b, Supplementary** Figure 4). Across replicates, specific A6 CDR3α and CDR3β clonotypes emerged with notable differences in peptide-binding profiles. All A6 variants bind minimally to non-Tax peptides and single-point Tax mutants of position 8, aligning with bulk NGS data (**Figure 4c**). The CDR3α variants NWD, DWD, and DWN, all with tryptophan at position 99 and either aspartic acid or asparagine at position 98 and 100, dominantly bind the peptide Tax P5Q (LLFG<u>Q</u>PVYV). In contrast with other TCR variants, these weakly bind the wild-type Tax peptide (LLFG<u>S</u>PVYV), and Tax P5W (LLFG<u>W</u>PVYV). Analogously, A6 CDR3β variants DGW, MGW, DGY, and MGY, all with a glycine at position 100, bind the Tax P5Q with a similar profile to the above noted CDR3α variants. A6 CDR3α variant DNW seems to bind Tax P5A (LLFG<u>A</u>PVYV) preferentially compared with other variants.

**Figure 5.**
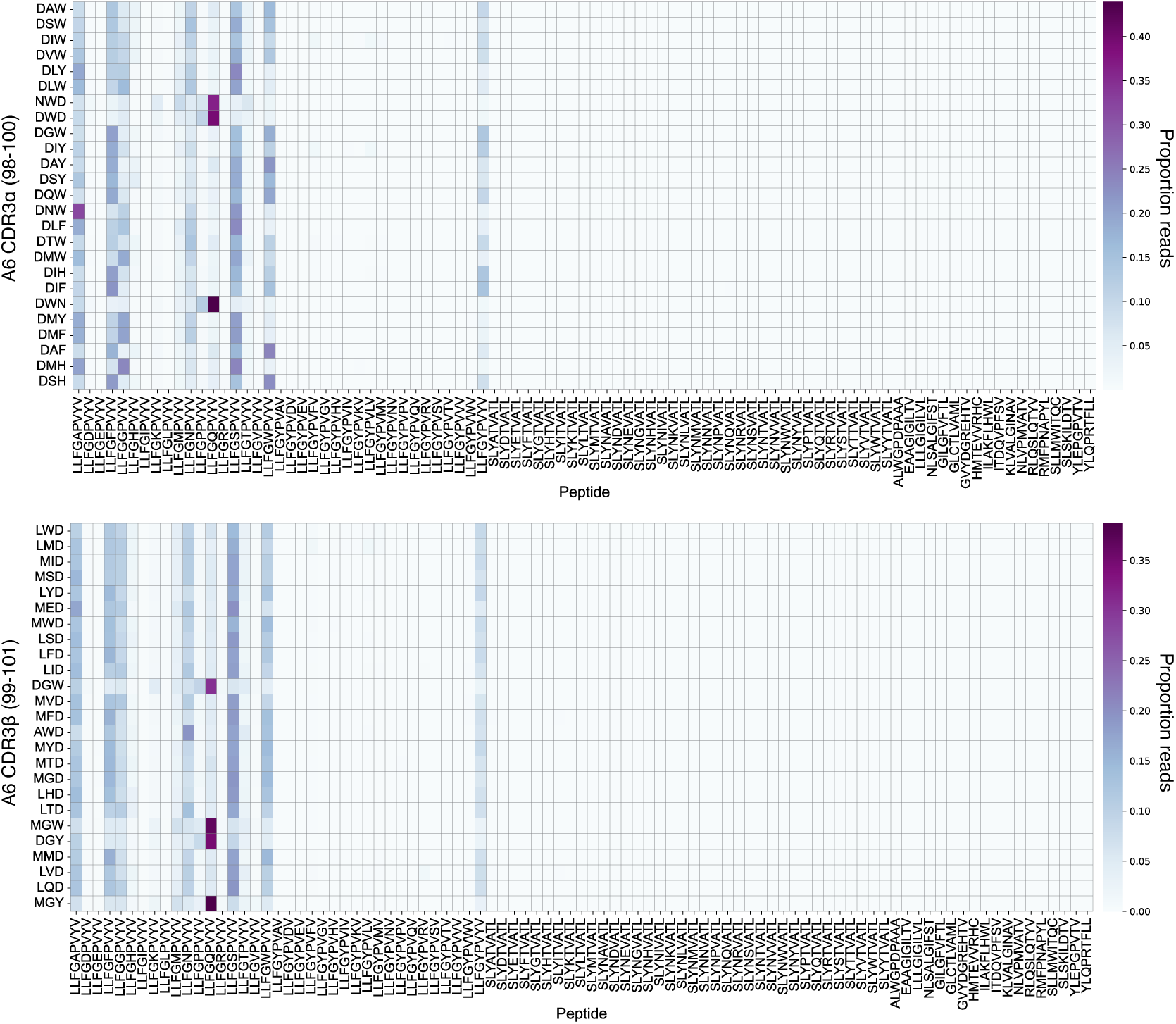
High-throughput sequencing of interacting TCR-pMHC pairs. Heatmaps depicting the proportion of reads for each A6 CDR3α- and CDR3β-peptide pair normalized to the total reads for each CDR3 variant. Data represents the mean of 3 technical replicates.

The 868 TCR variants exhibit less pronounced differences in peptide binding for SL9 variants, likely due to its higher affinity permitting reduced peptide stringency. Nonetheless, the 868 TCR shows apparent preferences for wild-type SLYNTVATL and, secondarily, SL9 P4A (SLY<u>A</u>TVATL) and SL9 P4I (SLY<u>I</u>TVATL) for both CDR3α and CDR3β (**Supplementary** Figure 4). Bulk NGS data in **Figure 4b** hints at 868 TCR variants binding the EBV epitope GLCTLVAML and several Tax epitopes. In single-cell data, we observe that the 868 CDR3α variant NSGP preferentially binds LLFG<u>N</u>PVYV over SL9 variants and weakly binds LLFG<u>H</u>PVYV. From bulk NGS of CDR3α- CDR3β amplicons, CDR3α NSGP does not exhibit strong preferences for a CDR3β sequence, pairing with TMSY as the top binder and the majority of the top 25 CDR3β sequences, explaining the apparent weak preference of all CDR3β sequences for LLFG<u>N</u>PVYV. CDR3α variant DSNY and CDR3β variant SMSF also weakly bind GLCTLVAML VLPs.

### Characterization of TCR clones identified in site-directed mutant library

To validate and characterize differences between enriched TCRs, we selected A6 CDR3α and CDR3β sequences with notably different Tax variant-binding profiles in the linked TCR-pMHC data (**Figure 5**). We leveraged the bulk sequencing dataset identifying enriched paired A6 CDR3α- CDR3β sequences to select A6 CDR3α and CDR3β pairs that are highly enriched by the pMHC- VLP library with differences in peptide (A6 CDR3α-CDR3β: DNW-MED, DWD-MGY, NWD- MGW, DLY-MED, DAW-AWD, DAW-LMD).

We established clonal yeast for the selected A6 TCR clonotypes and verified pMHC-VLP binding via two methods: staining with the 92-member pMHC-VLP library and staining with individual pMHC-VLPs for Tax mimotopes mutated at position 5 (LLFG<u>X</u>PVYV) (**Figure 6a, Supplementary** Figure 5). Although metrics of read proportion per VLP for NGS are not directly comparable to the proportion of clonal yeast stained with a given fluorescent VLP, the sequenced VLP barcodes generally align with individual VLP staining. For example, LLFG<u>F</u>PVYV VLPs bind A6 CDR3α-CDR3β variants DNW-MED, DLY-MED, DAW-AWD, and DAW-LMD but not NWD-MGW and DWD-MGY (**Figure 6a, Supplementary** Figure 5). Each of these TCRs binds single-point position 5 Tax epitopes with a starkly different profile. CDR3α/β variant DWD-MGY and NWD-MGW align closely to **Figure 5** data, exhibiting the strongest binding for LLFG<u>Q</u>PVYV and weaker but detectable binding for other position 5 mimotopes (K, M, N, P, T). Similarly, DNW-MED binds LLFG<u>A</u>PVYV, corresponding to **Figure 5** showing DNW as associated with this epitope. Peptide cross-reactivity can be changed significantly with just two altered residues, as observed between DAW-AWD and DAW-LMD. The general concordance between bulk sequencing of VLPs bound to individual TCR clonotypes (**Figure 4c**) and individual VLP staining confirms the accuracy of the sequencing approach in generating TCR-antigen association data. Collectively, these results demonstrate that single-residue mutations in CDR3α and CDR3β regions introduce substantial differences in TCR-antigen binding.

**Figure 6.**
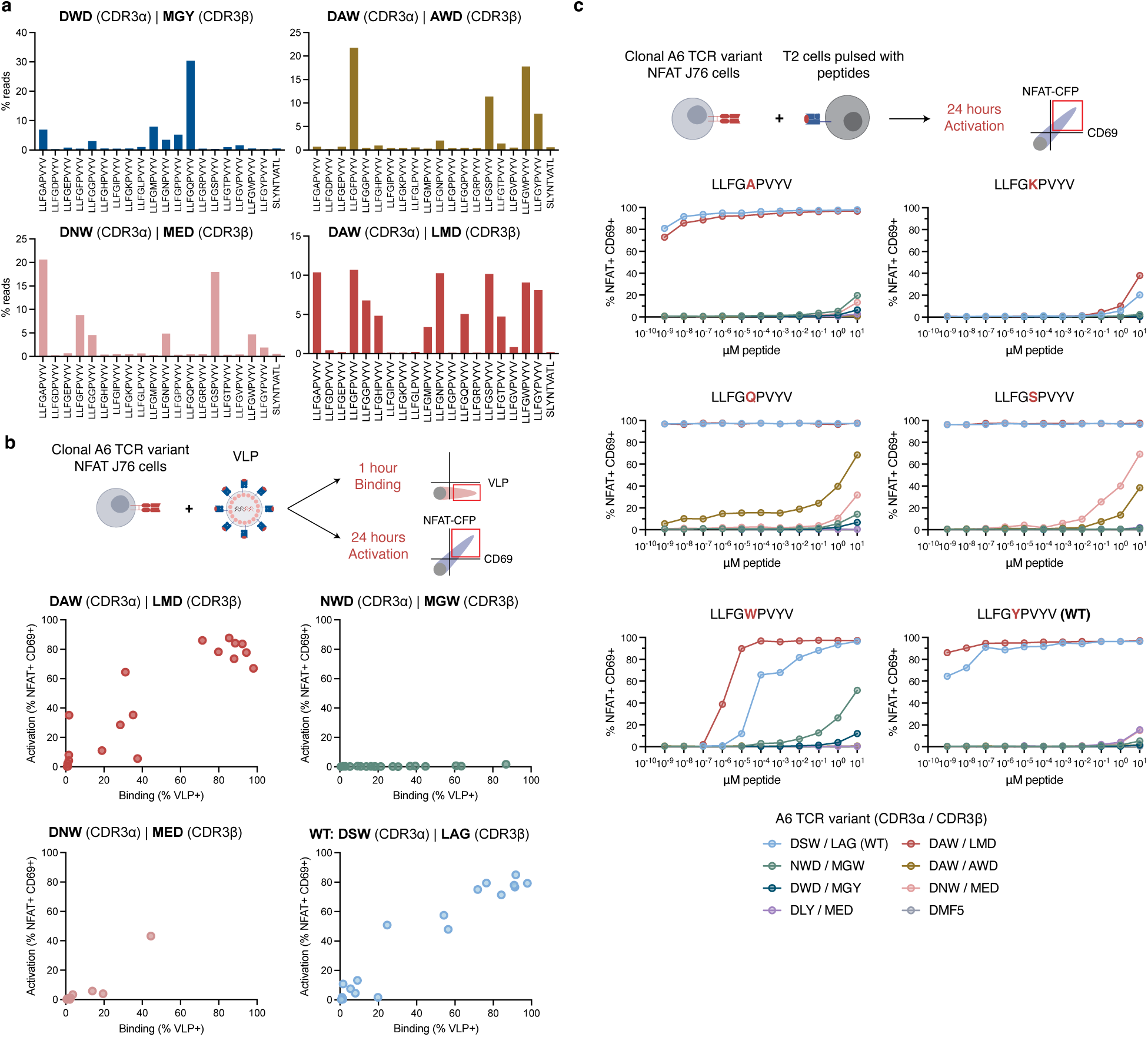
Validation of A6 TCRs with altered peptide binding in Jurkat J76 cells. a, Bar plot depicting proportion of reads per pMHC-VLP for four clonal A6 TCR variant-expressing yeast, focused on position 5 Tax peptide variants. **b,** Dot plots depicting proportion of clonal A6 TCR variant-expressing J76 cells bound (1 hour, %VLP^+^) and activated (24 hours, %NFAT^+^CD69^+^) by individual Tax variant pMHC-VLPs (position 5). **c,** Plots showing dose response of activation (%NFAT^+^CD69^+^) for clonal A6 variant J76 cells stimulated by peptide- pulsed T2 cells.

To examine how differences in our selections translate to T cell activity, we constructed clonal TCR-expressing cell lines for all selected A6 TCR variants using a TCR-null J76 Jurkat cell line expressing an NFAT-CFP reporter^63^ (**Figure 6b**). To examine binding and activation via pMHC- VLPs, we incubated cells with individual VLPs for one hour to evaluate binding and 24 hours to assess activation, detected as NFAT^+^ CD69^+^ cells. Although each TCR variant bound pMHC- VLPs, TCR variants displayed different sensitivities to activation. This data suggests pMHC binding is a necessary but insufficient condition for eliciting TCR-mediated activation, a phenomenon which has been previously observed in the context of pMHC multimers^64^. Although pMHC-VLP binding on TCR-expressing cells roughly aligns with yeast (i.e. A6 CDR3α-CDR3β variants NWD-MGW and DWD-MGY preferentially bind LLFG<u>Q</u>PVYV), significant differences in binding strength are apparent (i.e. DAW-LMD binds most epitopes strongly while DAW-AWD only weakly binds VLPs at this dose). This suggests that yeast, while a valuable tool for exploring TCR-pMHC binding, provide an incomplete picture of productive TCR-antigen engagement.

To extend this work, we examined TCR-mediated activation using peptide-pulsed T2 cells as APCs and across a range of antigen doses for select point mutant Tax peptides (**Figure 6c**). Aligning with VLP data, the wild-type A6 TCR and A6 CDR3α-CDR3β variant DAW-LMD potently respond to low doses for a range of peptides while the NWD-MGW and DWD-MGY variants fail to activate TCR-expressing cells. Single-residue substitutions in peptides and subtle changes to the CDR3 loops nonetheless yield measurably different antigen sensitivities, evidenced by the preferential response of DAW-AWD for LLFG<u>Q</u>PVYV compared with DNW-MED for LLFG<u>S</u>PVYV. The ability to comprehensively screen TCR–pMHC binding using pMHC-VLPs provides rich protein– protein interaction data that can uncover CDR3 variants with distinct peptide-binding profiles, even as questions remain about how factors like coreceptor engagement influence downstream T cell activation.

## Discussion

Efforts to understand natural immune recognition and engineer effective T cell therapies are constrained by a lack of scalable tools to systematically map the sequence determinants of TCR specificity. Here, we introduce VelociRAPTR, a high-throughput platform integrating yeast-displayed TCR libraries with pMHC-displaying virus-like particles and single-cell sequencing to rapidly screen millions of TCR-antigen interactions in a pooled format. Applying VelociRAPTR to interrogate A6 and 868 TCRs, we systematically mapped how combinatorial mutations to both CDR3α and CDR3β shape recognition to 92 pMHCs, including single-point mutants of target peptides. These large-scale, unbiased datasets reveal key features of TCR specificity and provide a foundation for deeper exploration of the TCR-pMHC binding interface.

Our results demonstrate that mutations to peptide-contacting residues in the CDR3 loops can sensitively restrict or expand TCR binding to related epitopes. While most CDR3 mutations are tolerated with only subtle shifts in specificity, several mutations notably alter recognition, as shown in the context of the A6 TCR. This nuanced profiling of specificity has important implications for therapeutic TCR development, where mitigating potentially lethal off-target reactivity is a major challenge^37–39^. Using VelociRAPTR as a multiplexed screening approach, we can simultaneously screen TCR variants against a target antigen while rigorously profiling potential cross-reactivity against a large panel of pMHCs early in development. This approach may enable more precise tuning of TCR specificity, improving the safety and efficacy of T cell therapies.

VelociRAPTR further enables scalable mapping of TCR-pMHC interactions as training data for deep learning models of TCR recognition. The traditional approach of screening yeast displayed libraries with recombinantly produced protein is slow, laborious, and low throughput. By eliminating recombinant protein production and using viral barcodes to capture pMHC identity, we can evaluate millions of interactions simultaneously. This allows us to generate rich TCR- pMHC binding data critical for predicting TCR recognition^40–42,58,60,62^.

By coupling the scale and robustness of yeast display with the ease of generating barcoded VLPs, VelociRAPTR enables pooled TCR-antigen mapping. This platform allows for rapid, quantitative profiling of the TCR sequence for altered binding of diverse peptide-MHCs, supporting safer and more precise engineering of therapeutic T cells and a valuable resource for deep learning models of TCR recognition. Finally, while our work focuses on TCR-pMHC interactions, we note that the fundamental principles of using yeast-VLP binding for profiling protein-protein interactions are not unique to this system. We anticipate that the strategy of coupling yeast display and VLPs is readily applicable to other membrane-associated proteins to generally map protein-protein interactions at unprecedented scale.

## Limitations

This platform offers numerous avenues for further improvement and exploration. While VelociRAPTR comprehensively maps TCR-antigen binding, our work corroborates the observation that pMHC binding to TCR can be insufficient for TCR activation. Integrating VelociRAPTR output with functional screens will bridge the gap between binding and activation thresholds. Although we deeply profile two TCRs in this work across an array ∼100 antigens, we believe that the fundamental signal-to-noise ratio of this approach would enable examination of a broader set of both TCRs and pMHCs, especially by incorporating recently described improved approaches for pooled VLP library production.

## Methods

### Media and cells

HEK293T cells (ATCC CRL-11268) were cultured in DMEM (ATCC) supplemented with 10% fetal bovine serum (FBS; Atlanta Biologics) and penicillin-streptomycin (Gibco). Jurkat-J76 cells, a Jurkat E6.1 subline without endogenous TCR alpha and beta chains, were a gift from M. Heemskerk^63^. J76 cells and T2 cells (ATCC CRL-1992) were cultured in RPMI-1640 (ATCC) supplemented with 10% FBS and penicillin-streptomycin. Yeast were grown to confluence at 30°C in SDCAA (pH 5) yeast media and sub-cultured into galactose-containing SGCAA (pH 5) media at an OD600 of 1 for two days at 20°C.

### Lentiviral particle production

VLPs were produced by transiently transfecting HEK293T cells with DNA and linear 25kDa polyethylenimine (PEI; Santa Cruz Biotechnology) or *Trans*IT-Lenti Transfection Reagent (Mirus Bio) at a 3:1 ratio of transfection reagent to DNA. HEK293T cells were seeded 24 hours prior to transfection at 8.33×10^4^ cells/cm^2^. DNA, transfection reagent, and Opti-MEM (ThermoFisher) were mixed and incubated for 10 minutes (TransIT-Lenti) or 15 minutes (PEI) at room temperature followed by dropwise addition to HEK293T cells. Following PEI transfection, medium was changed to complete DMEM with 25 mM HEPES 3-6 hours post-transfection.

VLP plasmid ratios consist of 1.9:1:1 of pHIV-pMHC plasmid to psPAX2.1 packaging plasmid (Addgene, plasmid #12260) to Gag-mCherry (Addgene, plasmid #85390). Total VLP plasmid amounts are 2.9 μg per well of a 6-well plate and 87 μg per T225 flask. Lentivirus plasmid ratios consist of 5.6:3:1 of pHIV transfer plasmid to psPAX2.1 packaging plasmid to pMD2.VSVG plasmid (Addgene, plasmid #12259). Total lentivirus plasmid amounts are 2.4 μg per well of 6- well plate and 72 μg per T225 flask. All pMHC single-chain trimer constructs were constructed with stabilizing disulfide bridge (Y84C in HLA-A*02-01 and L2C in peptide-β2M linker)^66,67^ and cloned into the pHIV backbone.

Lentivirus was collected at 48 and 72 hours and filtered through 0.45-μm polyethersulfone (Millex, Millipore Sigma), eliminating debris via centrifugation at 300g for 5 minutes prior to filtration for large volumes. Concentrated lentivirus (200×) was generated by ultracentrifugation at 100,000g for 45 minutes at 4°C . Supernatant was discarded, and the pellet was resuspended overnight in 100-130 μL of Opti-MEM at 4°C . Resuspended virus was aliquoted and stored at -80°C.

### Generation of pMHC-VLP library

LeAPS virus libraries were generated as previously described in Dobson et al^17^. Briefly, HEK293T cells were seeded 24 hours before transfection. A barcoded GFP-pLeAPS plasmid library with 18bp barcodes was constructed using randomized primers, followed by individual selection and validation of transformants. A total of 92 barcoded GFP-expressing pLeAPS plasmids and 92 pHIV-pMHC plasmids (Genscript) were used to produce VSVG-pseudotyped lentivirus in parallel within 24-well plates. After 48 hours, virus collections were performed, and HEK293T cells were seeded at 20% confluency. These cells were then transduced in duplicate 24-well plates with barcoded LeAPS and pMHC viruses (per well: 100 µL barcoded LeAPS virus and 700 µL pMHC virus), supplemented with 8 µg/mL diethylaminoethyl-dextran (Sigma-Aldrich) to enhance transduction.

After 48 hours, flow cytometry was used to assess transduction efficiency by detecting pMHC virus (mCherry^+^) and LeAPS-barcode virus (GFP+). The duplicate wells were pooled in proportion to the percentage of cells co-transduced with pMHC and LeAPS barcode viruses (mCherry^+^ GFP^+^) to maintain accurate representation of each library member. Pooled cells were then sorted (mCherry^+^ GFP^+^) to establish the pMHC-VLP-packaging cell line. To produce pMHC-VLPs, packaging cells were seeded following the same protocol as HEK293T cells and transfected with psPAX2.1 and Gag-mCherry plasmids using the TransIt-Lenti (Mirus Bio) transfection reagent.

Library pMHC-VLPs were harvested and concentrated at 48 and 72 hours, following the previously described procedure.

### Generation of TCR-expressing yeast libraries

#### Yeast-displayed scTv design

The A6 TCR was expressed as a single-chain TCR (scTv) consisting of two linked variable regions (VH and VL), optimized for yeast display as previously described^26^. The alpha and beta variable domains are connected by a flexible linker in the orientation Vβ-L-Vα at the C-terminus of the yeast cell wall protein Aga2. scTv constructs were expressed from the yeast display vector pCT302 with an Aga2 leader peptide sequence and N- and C-terminal expression tags, hemagglutinin (HA) and Flag epitope, respectively^68,69^.

Randomized CDR3α/β yeast libraries were generated by overlap extension PCR of scTv fragments introducing diversity with primers encoding NNK degenerate codons. To ensure expression of only randomized scTvs, the template vector and fragments encoded stop codons at all mutagenized positions in both CDR3α/β. Randomized scTv PCR products and linearized pCT302 vector were mixed at a mass ratio of 5:1 and electroporated into electrocompetent RJY100 yeast^70^. The final scTv libraries contained a minimum of 10× coverage of the theoretical library diversity. The A6 TCR-CDR3β library consisted of mutated CDR3α positions D98, S99, and W100. The A6 TCR- CDR3α/β library consisted of mutated CDR3α positions D98, S99, and W100, and CDR3β positions L99, A100, and G101.

### Generation of monoclonal TCR yeast

Individual TCRs were assembled into pCT302 vectors as single-chain TCRs with selected CDR3α and CDR3β sequences. Constructs were electroporated into electrically competent RJY100 yeast as described above^70^.

### pMHC monomer production

Recombinant soluble HLA-A*02:01 single-chain trimer was produced using a baculovirus expression system with High Five (Hi5) insect cells (ThermoFisher). The Tax-A2 construct was cloned into a pVL1393 insect cell expression vector in the format of signal peptide, peptide, β2M, HLA-A*02-01, an AviTag biotinylation site (GLNDIFEAQKIEWHE), and a poly-histidine (8× His) purification site, connected via flexible Gly-Ser linkers. Plasmid DNA was transfected into SF9 insect cells with BestBac2.0 DNA (Expression Systems) with Cellfectin II (Thermo Fisher) to generate baculovirus. Hi5 insect cells were transduced and incubated for 48-72 hours. Secreted protein was purified from pre-conditioned media supernatant with Ni-NTA resin and via size exclusion chromatography using an S200 increase column on an AKTAPURE FPLC (GE Healthcare).

### Recombinant pMHC yeast selection

Yeast selections were completed with streptavidin-coated magnetic beads (Miltenyi Biotec) coupled to biotinylated Tax-A2 monomer, confirmed by streptavidin gel shift assay^71^. Three iterative rounds of selection were completed, culturing and inducing yeast after each round. Each round included a negative selection step with uncoated streptavidin beads to clear non-specific yeast.

### Individual pMHC-VLP yeast staining

Monoclonal TCR-expressing yeast were stained in FACS buffer (PBS, 0.1% BSA, 1 mM EDTA) with an anti-Flag epitope antibody (clone R5, Biolegend) in FACS buffer (1:100) for 10 minutes, washed once, and assessed for expression induction via flow cytometry. Flag-stained yeast were incubated for 1 hour at 4C in a total staining volume of 50 μL FACS buffer with a specified amount of concentrated VLP or 100 μL consisting of 50 μL unconcentrated VLPs and FACS buffer. VLP- stained yeast were washed 2× in FACS buffer prior to analysis via flow cytometry. If a secondary stain is used, cells were stained with an anti-β2M antibody (clone 2M2, Biolegend) in FACS buffer (1:100) at 4°C for 20 minutes.

### pMHC-VLP library screening of TCR yeast libraries

Both magnetic- and flow cytometry-based enrichments were used to select pMHC-binding yeast following conventional yeast display protocols^6^. Naïve libraries were screened at a minimum of 10-fold coverage of the estimated library diversity for the first selection round (470 million per selection for an estimated CDR3α/β library diversity of 4.7×10^7^ and 15 million per selection for an estimated CDR3β library diversity of 7.0×10^3^). Subsequent rounds were executed on 25 million and 10 million yeast per round, respectively.

For flow cytometry-based selections (FACS), TCR-expressing yeast were incubated with single pMHC VLPs (0.5 μL concentrated VLP and 49.5 μL FACS buffer per 20 million yeast) or a library of pMHC VLPs (50 μL concentrated VLP and 20 μL FACS buffer per 20 million yeast) rotating at 4°C for 1 hour. Yeast were washed 3× in FACS buffer and sorted on fluorescence (mCherry^+^).

For magnetic-activated cell sorting (MACS) selections, yeast were negatively selected to eliminate non-specific binders by incubation with anti-PE magnetic microbeads (Miltenyi Biotec). Yeast were subsequently stained with VLPs as described above, followed by incubation with PE anti- β2M antibody (clone 2M2, Biolegend) in FACS buffer (1:50) at 4°C for 20 minutes, 3 or more washes, and incubation with anti-PE magnetic microbeads (1:100, v/v) rotating at 4°C for 2 hours. Between rounds of selection, yeast were grown to confluence and induced as described above.

### Deep sequencing of scTv and pMHC-VLP libraries

Pooled plasmids from each round of selection were isolated using a Zymoprep II Yeast Miniprep kit (ZymoResearch). Amplicons capturing both CDR3α and CDR3β were generated using purified miniprep plasmids via 30 cycles of PCR. The amplicons were submitted to Genewiz for Amplicon-EZ NGS analysis, generating paired 2 × 150 nt reads. Data was processed to extract peptide sequences between constant flanking regions.

To extract bulk pMHC binding data, 5 million enriched yeast were stained with the VLP library as described, washed 4× to remove unbound VLPs, and treated with 2U (0.5 μL) of Zymolyase (Zymo Research) in 40 μL of nuclease-free water for 20 minutes at room temperature to strip bound VLPs. Yeast were pelleted at 5000g for 1 minute and supernatant was extracted as a template for RT-PCR. RT-PCR was executed using the TaqMan Fast Virus 1-Step Master Mix (ThermoFisher) (25 cycles) to add partial Illumina Truseq adapters prior to sequencing via Amplicon-EZ NGS analysis by Genewiz. The VLP library was sequenced directly using the TaqMan Fast Virus 1-Step Master Mix (1μL of VLP).

### CDR3-pMHC sequencing

Enriched yeast were stained with the 92 pMHC-VLP library as described, washed 4× to remove unbound VLPs, then single-cell sequencing was performed and the resulting libraries were sequenced with Element’s Aviti platform.

### Functional validation in monoclonal TCR lines

CD8^+^ J76 cells were modified to express a fluorescent reporter of nuclear factor of activated T cells (NFAT)^63^. NFAT-CFP-encoding retrovirus was generated by transfecting HEK293T cells with pSIRV-NFAT-CFP (constructed from pSIRV-NFAT-GFP, Addgene, plasmid #118031), pUMVC (Addgene, plasmid #8449), and pMD2.VSVG as described above. CD8^+^ J76 cells were transduced by unconcentrated NFAT-CFP-encoding lentivirus, sorted as single cells into a 96-well plate, cultured as described for 3 weeks. Half of each clonal population was stimulated with phorbol myristate acetate (PMA,10 ng/mL, ThermoFisher) and ionomycin (1 μg/mL, ThermoFisher). Reporter CFP expression was compared between stimulated cells and unstimulated cells and clones with the highest differential were selected. The selected clone is henceforth referred to as NFAT-CFP CD8^+^ J76.

Lentiviral TCR constructs were formatted as TCRβ-P2A-TCRα and cloned into pHIV backbone vectors. NFAT-CFP CD8^+^ J76 cells were transduced with unconcentrated TCR lentivirus generated as described above (1 mL per 1M cells), TCR expression was quantified by flow cytometry (anti-TCR antibody, clone IP26, Biolegend), and TCR^+^ cells were sorted on a FACSAria cell sorter. TAP-deficient T2 cells were pulsed with individual peptides at a range of doses for 4 hours. 100,000 peptide-pulsed T2 cells were incubated with 100,000 TCR^+^ NFAT-CFP J76 cells overnight for each condition. Cells were washed and stained with anti-CD69 (Biolegend, clone FN50) and anti-CD19 (Biolegend, clone HIB19) in FACS buffer, both at a 1:200 dilution. Alternatively, cells were stained with unconcentrated pMHC-VLP for 1 hour at 4°C or overnight at 37°C. Cells were analyzed for binding (1 hour) or activation (overnight) on a Cytoflex S flow cytometer.

## RESOURCE AVAILABILITY

### Lead contact

Further information and requests for resources and reagents should be directed to and will be fulfilled by the lead contact, Michael Birnbaum.

### Materials availability

Plasmids are available upon request.

### Data and code availability

Data reported in this paper is available from the lead contact upon request.

## Acknowledgements

We thank the Koch Institute’s Robert A. Swanson (1969) Biotechnology Center for their technical support, especially the Flow Cytometry Facility and MIT BioMicro Center. We thank S. Levine, N. Kamelamela, and G. Paradis for helpful discussions and suggestions. This work was supported in part by the Koch Institute Frontier Research Program through the Michael (1957) and Inara Erdei Fund and the Casey and Family Foundation Research Fund, the Packard Foundation, NIH Director’s New Innovator Award (DP2-AI158126), U.S. Army Medical Research (W81XWH2210300), and Pfizer Inc. to M.E.B.; the National Institutes of Health (R01CA218094) to D.G., and a Canadian Institutes for Health Research Doctoral Foreign Study award to S.A.G. This work was delivered as part of the MATCHMAKERS team, of which M.E.B. is a member, supported by the Cancer Grand Challenges partnership financed by CRUK (CGCATF- 2023/100001), the National Cancer Institute (OT2CA297463), and The Mark Foundation for Cancer Research. This content is solely the responsibility of the authors and does not necessarily represent the official views of the National Institutes of Health.

## Author contributions

Conceptualization, S.A.G., J.K., F.K. and M.E.B; methodology and experiments, all authors; data analysis, S.A.G; writing, S.A.G. and M.E.B.; review and editing, all authors.

