## Supplemental Information for "Deep mapping of the TCR-antigen interface using pMHC-pseudotyped viruses and yeast display"

### Supplementary Figures

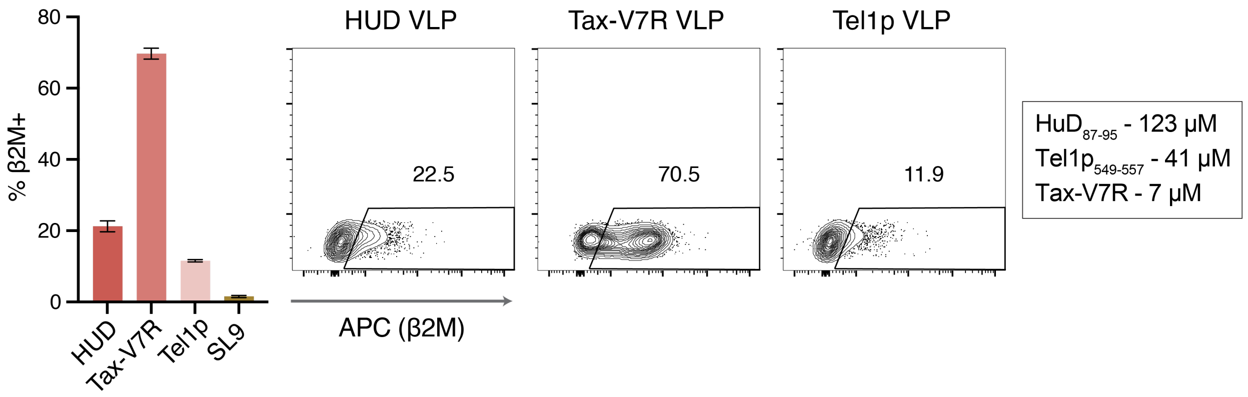


**Supplementary Figure 1.** Bar plot depicting the proportion of A6 yeast stained with a A6-binding pMHC-VLPs (β2M$^{+}$) for peptides of varying affinities. Representative flow plots are shown. Bar plot data represents the mean of 3 technical replicates ± S.D.

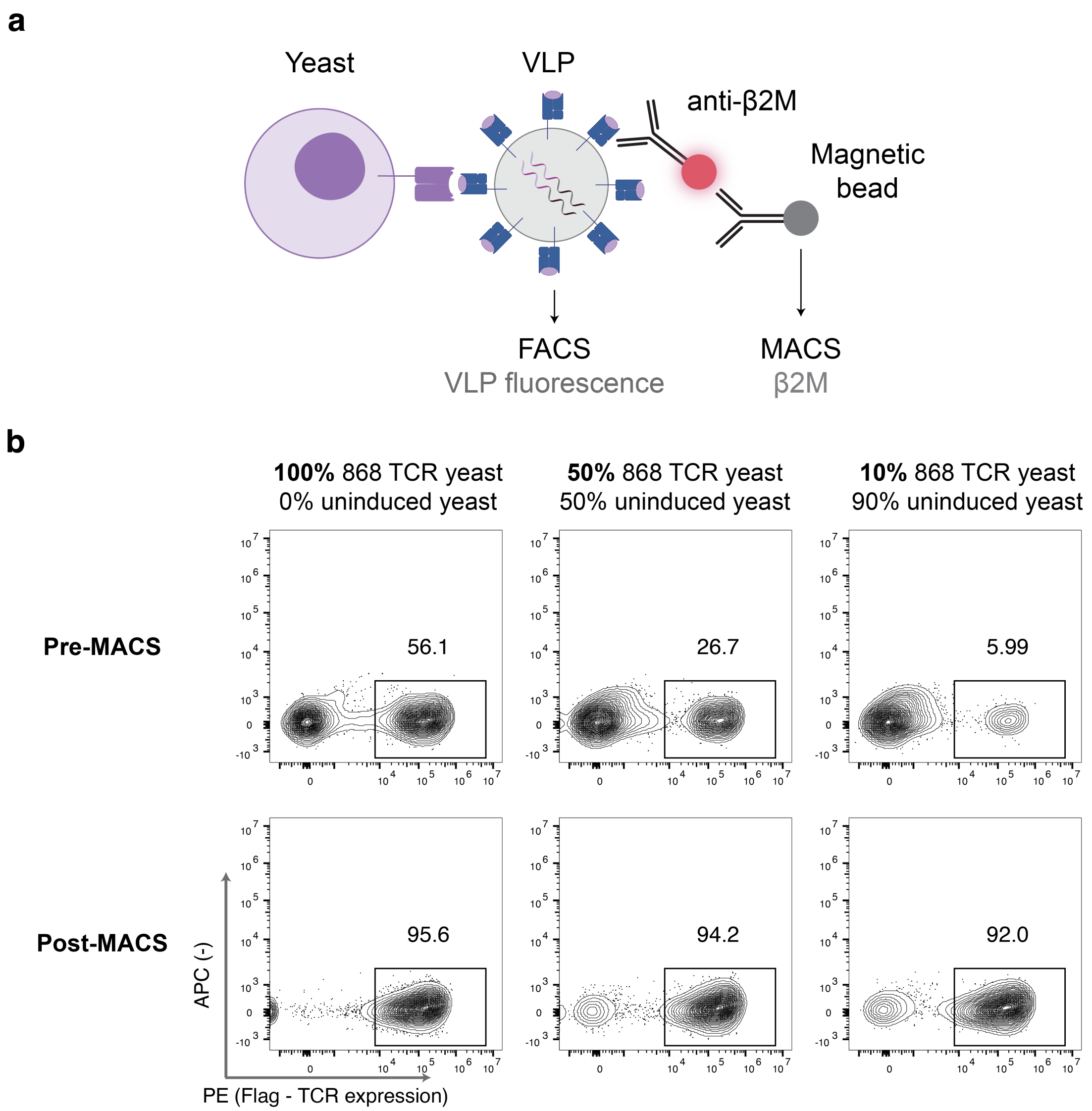


**Supplementary Figure 2. a,** Schematic illustrating both fluorescent- and magnetic-based cell sorting to isolate VLP-bound yeast. **b,** On- and off-target yeast (wild-type TCR-expressing and uninduced, respectively) were mixed at different ratios, stained with SL9 pMHC-VLPs, and purified via MACS to assess the efficiency of using MACS to purify VLP-bound yeast. Plots depict the proportion of TCR-expressing yeast before and after MACS.

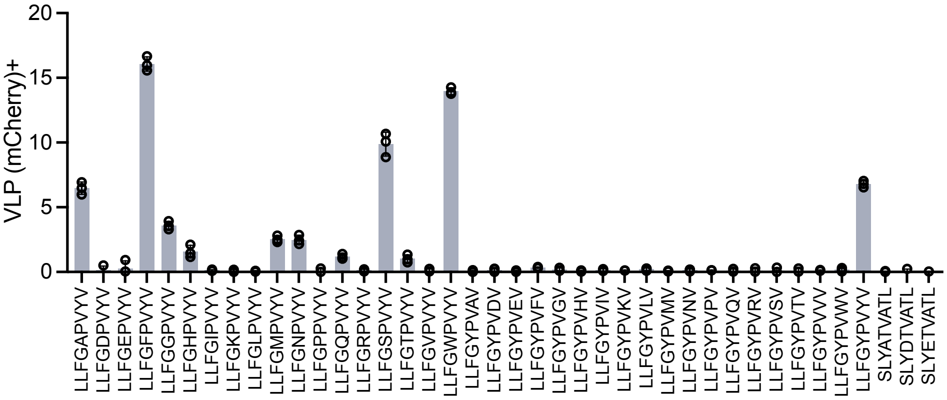


**Supplementary Figure 3.** A6 TCR yeast, varied at both CDR3α and CDR3β as described, were stained individually with unconcentrated pMHC-displaying VLPs. Bar plot depicts the proportion of all yeast identified via flow cytometry as $VLP^+$.

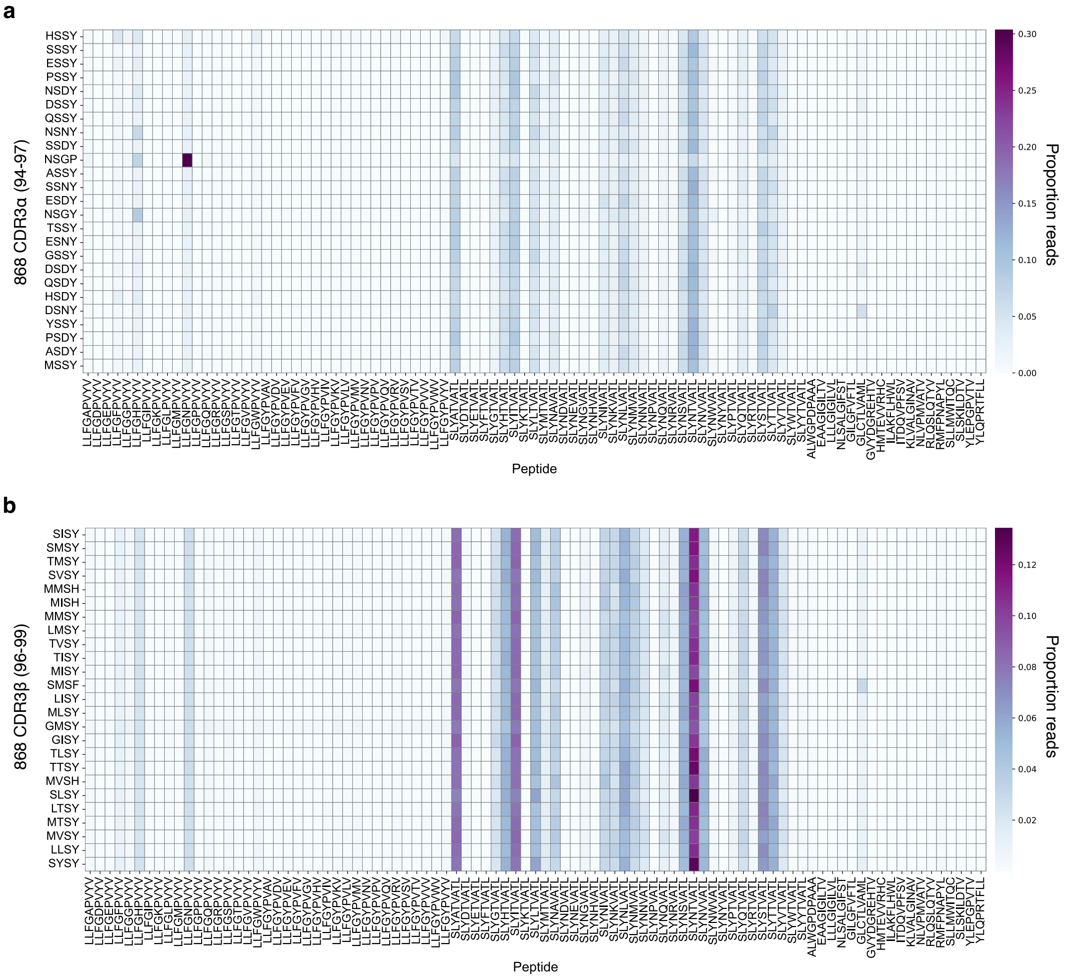


**Supplementary Figure 4.** Heatmaps depicting the proportion of reads for each 868 CDR3α- and CDR3β-peptide pair normalized to the total reads for each CDR3 variant. Data represents the mean of 3 technical replicates.

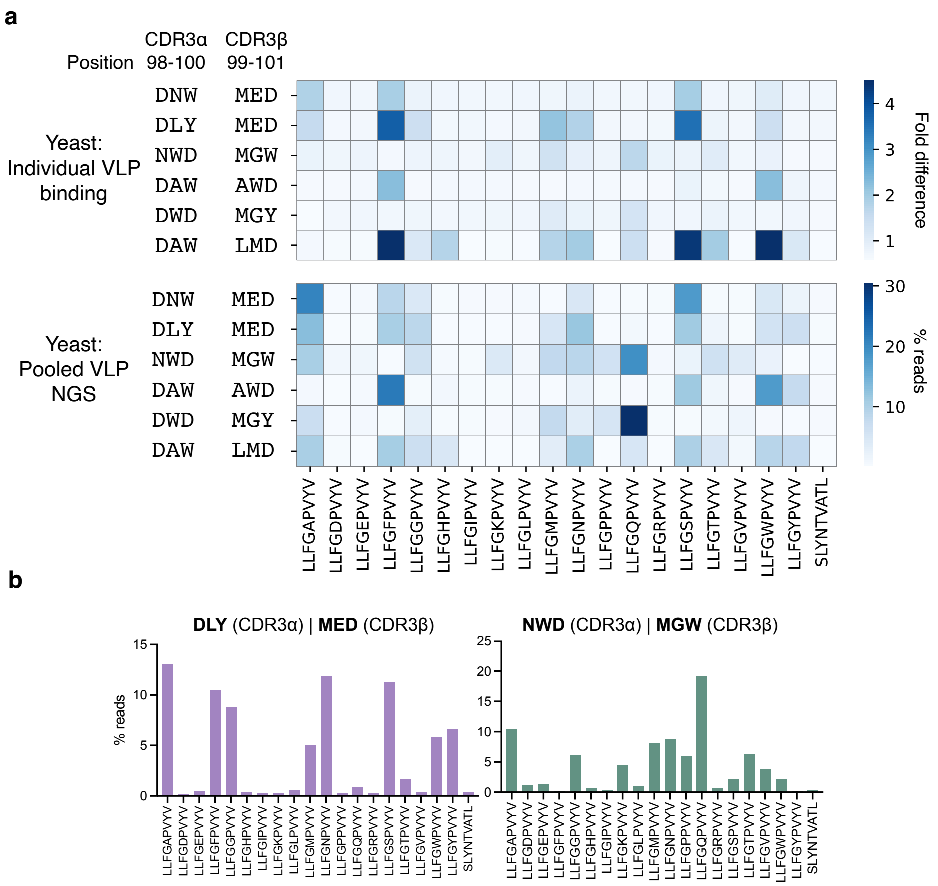


**Supplementary Figure 5. a**, Individual A6 TCR variants were expressed on yeast and stained with a 92-member pMHC-VLP library followed by NGS of bound pMHC-VLPs to reveal Tax epitope-binding profile of each variant. Reads per epitope are normalized to the total number of reads per variant. Each CDR3α-CDR3β clonotype is represented by three CDR3α residues and three CDR3β residues (i.e. DNW-MED: DNW = CDR3α and MED = CDR3β). Individual A6 TCR variants were stained with individual A6 pMHC-VLPs. Fold difference represents ratio of $Flag^{+}\beta2M^{+}$ yeast to $Flag^{-}\beta2M^{+}$ yeast (signal-to-noise ratio). Data represents the mean of two technical replicates. **b,** Depiction of heatmap data analogous to Figure 5 for A6 CDR3α-CDR3β clonotypes DLY-MED and NWD-MGW, showing the proportion of reads for position 5 Tax variants and a negative control.

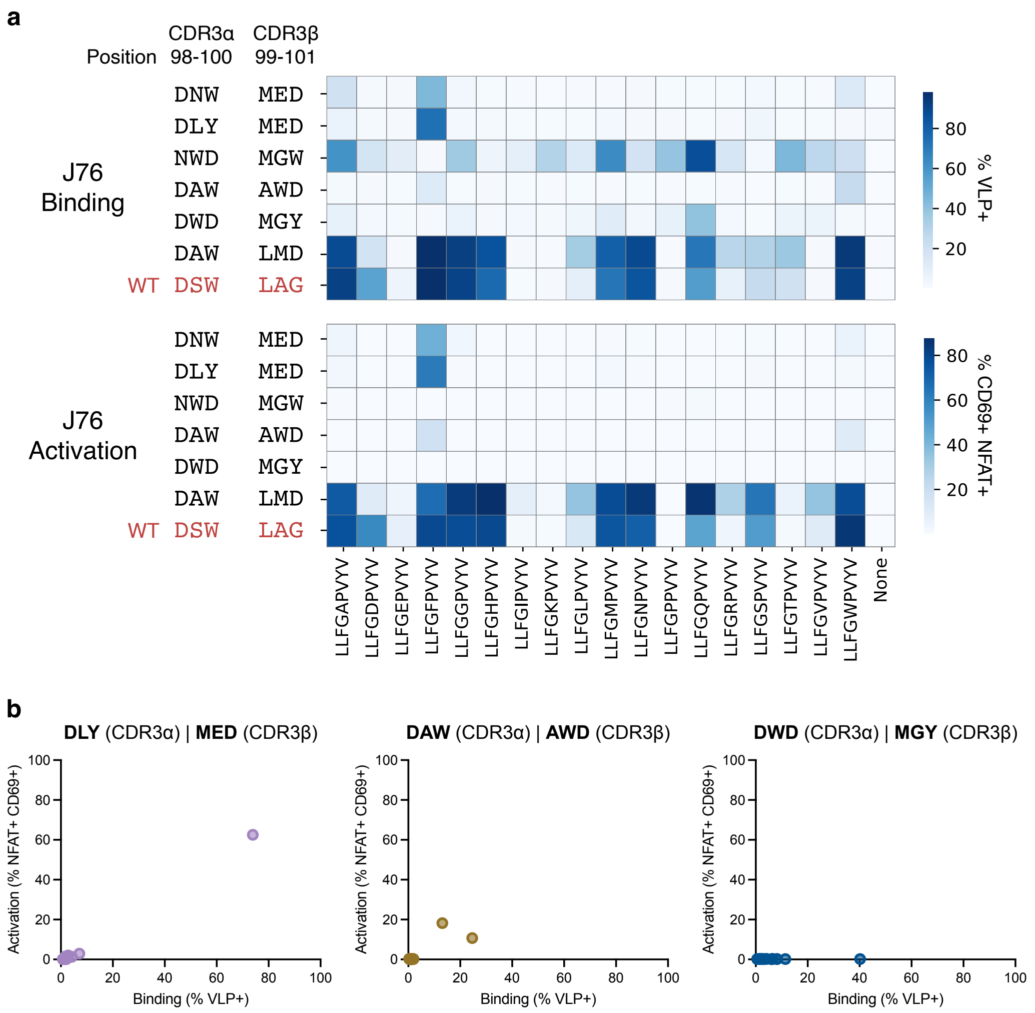


**Supplementary Figure 6. a,** Heatmap depicting binding (% VLP$^+$) and activation (% NFAT$^+$CD69$^+$) of clonal A6 TCR variants by individual Tax variant pMHC-VLPs. **b,** Dot plots depicting correlation between activation and binding by individual Tax variant pMHC-VLPs for select clonal A6 TCR variants not presented in Figure 6.
